# Deep autoregressive generative models capture the intrinsics embedded in T-cell receptor repertoires

**DOI:** 10.1101/2022.09.01.505405

**Authors:** Yuepeng Jiang, Shuai Cheng Li

## Abstract

T-cell receptors (TCRs) play an essential role in the adaptive immune system. Probabilistic models for TCR repertoires can help decipher the underlying complex sequence patterns and provide novel insights into understanding the adaptive immune system. In this work, we develop TCRpeg, a deep autoregressive generative model to unravel the sequence patterns of TCR repertoires. TCRpeg outperforms state-of-the-art methods in estimating the probability distribution of a TCR repertoire, boosting the accuracy from 0.672 to 0.906 measured by the Pearson correlation coefficient. Furthermore, with promising performance in probability inference, TCRpeg improves on a range of TCR-related tasks: revealing TCR repertoire-level discrepancies, classifying antigen-specific TCRs, validating previously discovered TCR motifs, generating novel TCRs, and augmenting TCR data. Our results and analysis highlight the flexibility and capacity of TCRpeg to extract TCR sequence information, providing a novel approach to decipher complex immunogenomic repertoires.

## Introduction

The adaptive immune system consists of highly diverse B and T-cells whose unique receptors can recognize enormous pathogens in vertebrates. The generation of these highly diverse receptors arises mainly from the genetic recombination of DNA segments from V, D, and J genes through V(D)J recombinations^1, 2^. T-cells play an essential role in antiviral defense by selectively eliminating virus-infected cells^3^. Their ability to recognize specific short peptides; that is, peptide antigens bound to the major histocompatibility complex (MHC) molecules are determined primarily by their unique receptor proteins^4, 5^. A receptor contains an *α* polypeptide chain and an *β* polypeptide chain, both of which consist of two extracellular domains: the variable (V) region and the constant (C) region^6^. The variable regions of the TCR *α* - and *β* - chains both have three complementarity-determining regions (CDRs) that contribute to the specificity of antigen recognition. Among these CDRs, the CDR3 region of the TCR *β* chain plays a pivotal role in the recognition of the peptides presented by MHC. In contrast, the CDR1 and CDR2 regions contribute minor effects to direct antigen recognition^6, 7^. Due to the importance of the highly diverse CDR3 region of the TCR *β* chain in antigen recognition and data availability, this work focuses on deciphering the underlying pattern of the CDR3 sequence.

Advancement in high-throughput sequencing techniques of the T-cell receptor repertoire provides a census of T-cells found in blood or tissue samples^8–11^. Large-scale sequencing data promote the investigation of the composition of immune repertoires, characterizing adaptive immune responses, and developing descriptive models. The sampled repertoire of TCR serves as an indicator of the complete repertoire, reflecting the pathogenic history or the immune response to stimuli^12–15^, with clinical applications including cancer prediction and anticipation of immunotherapy. For example, Han *et al*. developed a statistical index named TIR index based on TCR to predict response and survival outcomes after immunotherapy^16^. Beshnova *et al*. defined a cancer score for a given patient based on the predictive model trained on specific TCR sequences that are assumed to be simply associated with cancers^17^.

Despite the success in predictive tasks associated with T-cell repertoires, precise probabilistic distribution modeling is demanding. Given that TCR repertoires possess extremely large diversity, the sampled repertoires from different samples, or even from the same donors will often differ significantly. Consequently, characterizing the sequence pattern of a given repertoire from a probabilistic manner is more reliable than modeling with raw TCR sequences and read counts, with many potential applications such as estimating the relative ratio of CD4^+^ to CD8^+ 18, 19^ and investigating the differences in sequence characteristics between functional T-cell subsets^20, 21^. Conventionally, modeling the sequence pattern behind a TCR repertoire is disentangled into two processes: generation (V(D)J recombination)^22, 23^ and selection^19, 24, 25^. The ultimate probability assigned to a TCR sequence is the product of the selection factor and the generation probability inferred from the selection process and the generation process, respectively. However, the generation models learned from different individuals share a high mutual similarity^22, 25^, indicating that the selection process plays a central role in discriminating the TCR repertoires sampled from different individuals. Therefore, instead of two-step disentanglement, we can infer the probability of TCR sequences end-to-end.

In this work, we introduce a new probabilistic model, TCRpeg, that utilizes deep learning techniques to learn the underlying sequence patterns of TCR repertoires. Specifically, TCRpeg employs the architecture of the deep autoregressive model with gated recurring units (GRU)^26^ layers to characterize the repertoire through the flexible and non-linear structure of deep neural networks. TCRpeg can infer the sequence probability distribution with higher accuracy than other probabilistic models, boosting the performance from 0.672 to 0.906 measured by the Pearson correlation coefficient. We then applied the model to profile TCR subrepertoires and found that a simple probabilistic classifier can achieve high predictive performance. TCRpeg also provides high-quality latent vector representations for TCR sequences. Based on these vector encodings of TCR sequences, we built a fully connected neural network to classify the cancer-associated TCRs and SARS-CoV-2 epitope-specific TCRs, achieving 0.844 and 0.872 AUC, respectively; higher than DeepCAT^17^’s AUC 0.768 but slightly lower than TCRGP^27^’s AUC 0.882. As a generative model, TCRpeg can generate new TCR sequences, among which more than 50% share the same antigen specificity as the sequences used in training according to the TCRMatch^28^ with a scoring threshold of 0.90, while the other two generative models, TCRvae^29^ and soNNia^19^, achieve a proportion of less than 40%. Further, TCRpeg helps data augmentation; it shows a 7.4% accuracy gain in predicting cancer-associated TCRs using the DeepCAT^17^ model.

## Results

### Autoregressive generative model for TCR sequences

Previously, the probabilistic sequence pattern of a TCR repertoire was modeled by the two disentangled processes of generation^22, 23^ and selection^19, 24, 25^ (e.g., soNNia^19^) or the variational autoencoder with convolutional neural networks (CNNs) as encoder and decoder (TCRvae^29^). Although both models achieved satisfactory performance, they lack the elegance to handle variable-length TCR sequence data. The two types of models pad each sequence to a fixed length with an extra token representing the padding positions. However, the introduction of the extra token could introduce noise to the original data and partially conceal useful information about the diversity of sequence lengths, which is important for antigen specificity^30, 31^.

In the past decade, deep learning models have achieved considerable success in handling sequential data ^26,32–35^. An autoregressive model processes the sequential data using observations from previous stages to infer the entry at the next time point. In the context of the TCR sequence, we can apply an autoregressive model to infer a residue using the amino acid subsequence proceeding from it. Therefore, we built TCRpeg, an autoregressive model that formulates the probability of a TCR sequence ***x*** as *p*(***x***|***θ***), where the parameters ***θ*** capture the latent evolutionary patterns to generate ***x***. The probability density *p*(***x|θ***) can be calculated by the product of probabilities conditioned on previous residues along a sequence with length *L* through an autoregressive likelihood

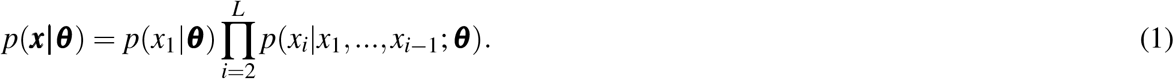

Figure 1 shows the TCRpeg workflow. We utilized gated recurrent units (GRUs)^26^, commonly adopted in recurrent neural networks, to model the autoregressive likelihood (Methods). Recurrent neural network models might encounter a gradient explosion for long peptide sequences^36, 37^. However, TCR sequences contain mainly 12 to 17 residues (Supplementary S1). Thus, we can parameterize the generative process with feed-forward GRU models that aggregate dependencies in sequences through the transmitting hidden features controlled by the gate functions.

**Figure 1.**
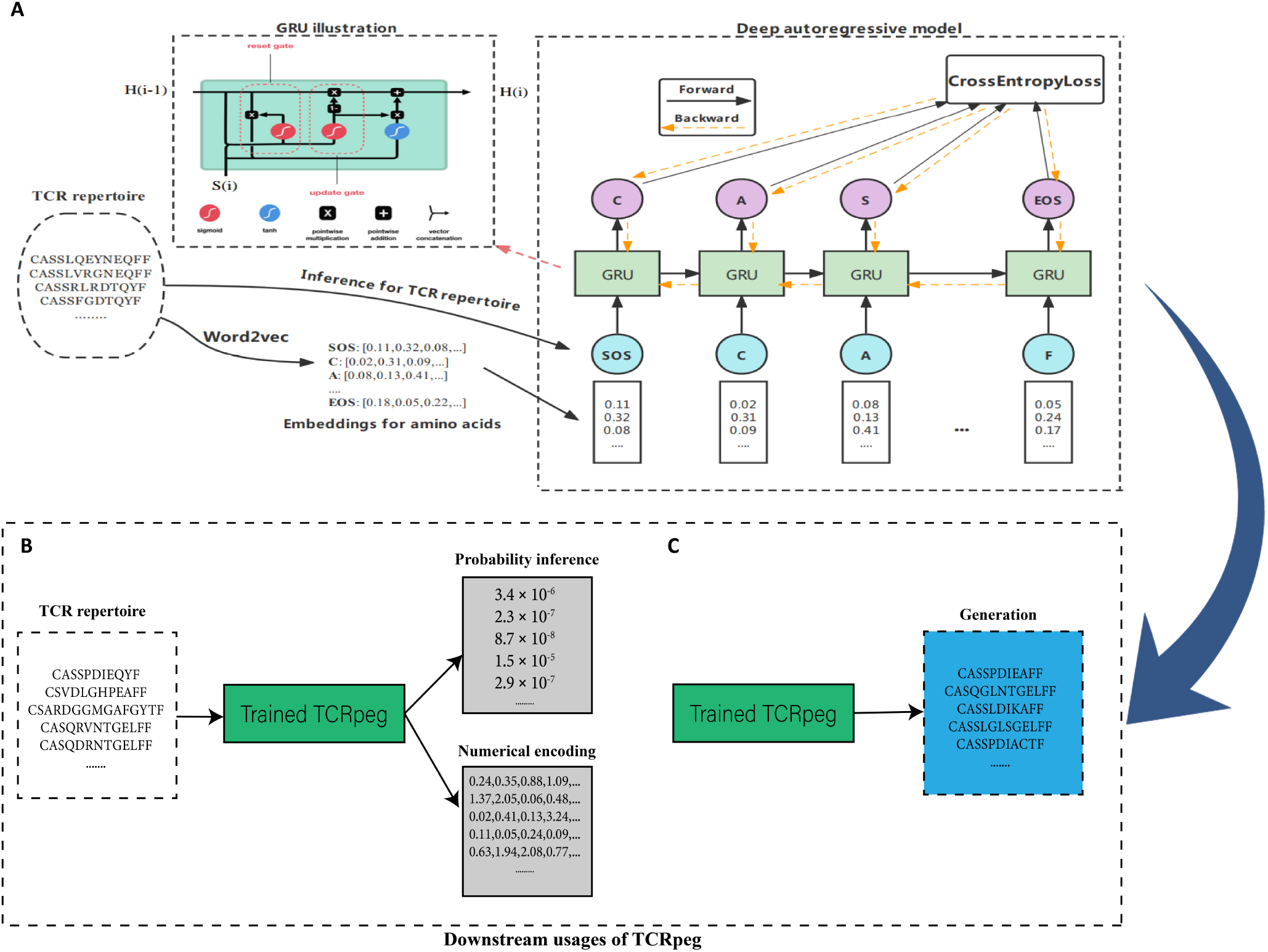
Workflow of TCRpeg to infer probabilistic patterns of immune receptor repertoires. (**A**) We have implemented a deep autoregressive network with GRU layers to process TCR sequences of different lengths to learn the hidden sequence pattern. The word2vec algorithm is first applied to the TCR repertoire to learn the numerical representations of each amino acid, regarding amino acids and TCR sequences as “Words” and “Sentences”. Then the TCR sequence is inputted into the deep autoregressive model sequentially. The model is updated by the gradient descent algorithm with the cross-entropy loss between the output logits and true labels. The trained TCRpeg model can be readily extended to downstream usages, including probability inference, encoding TCRs (**B**), and generating similar new TCRs (**C**). These functions and applications of TCRpeg are further elaborated in the Results section.

Training a GRU model requires vector representations for each amino acid. Instead of using one-hot encodings or predefined characteristics of the analysis of principal components in biochemical features^17^, we adopted the word2vec algorithm^38^ to adaptively learn the embeddings for each amino acid from the TCR sequencing data by treating an amino acid as a “Word” and each TCR sequence as a “Sentence” (Method). Then, TCRpeg can be trained in a forward language modeling manner. To estimate the probability of a given TCR sequence, we applied Eq.1 to the pre-trained TCRpeg. Details of the architecture of TCRpeg, the training, and inferring processes are included in the Methods.

### TCRpeg infers functional TCR repertoire probability distribution

First, we evaluated the probability distribution of the TCR sequences inferred by TCRpeg and compared its accuracy with the other two probabilistic models, soNNia^19^ and TCRvae^29^. To assess and compare their performance, we constructed a universal TCR repertoire from a large cohort of 743 individuals from Emerson *et al*.^39^, following a similar data preprocessing strategy in Isacchini *et al*.^19^. Specifically, we pooled the unique nucleotide sequences of TCRs from all individuals and constructed a universal TCR repertoire. The universal repertoire was randomly divided into training and testing subrepertoires by a 50:50 split to ensure consistency with soNNia^19^ and TCRvae^29^. Then we trained TCRpeg, soNNia, and TCRvae on the training set (Methods).

We evaluated the three models, each to estimate a probability distribution *P_in f er_*(***x***) for the test set; TCRpeg shows high accuracy with substantial improvement over soNNia and TCRvae, but requires lower resources to train. Prediction accuracy can be quantified using the Pearson correlation coefficient *r* between the inferred and true probability distributions, i.e., *P_in f er_*(***x***) and *P_data_*(***x***), on the test set (Methods). TCRpeg achieved *r* ≃ 0.906; however, soNNia and TCRvae obtained *r* ≃ 0.672 and *r* ≃ 0.653, respectively (Fig. 2A-2C). TCRpeg also performs stably and robustly when training on a small proportion of training data consisting of only 2 × 10^5^ TCR sequences (Supplementary S2). In addition to the substantial accuracy improvement, TCRpeg converges faster and costs significantly less GPU memory (Fig. 2D-2F). It converges within five epochs, whereas the other two methods require around 30 epochs. Moreover, SoNNia and TCRvae consume six times and three times more memory than TRCpeg.

**Figure 2.**
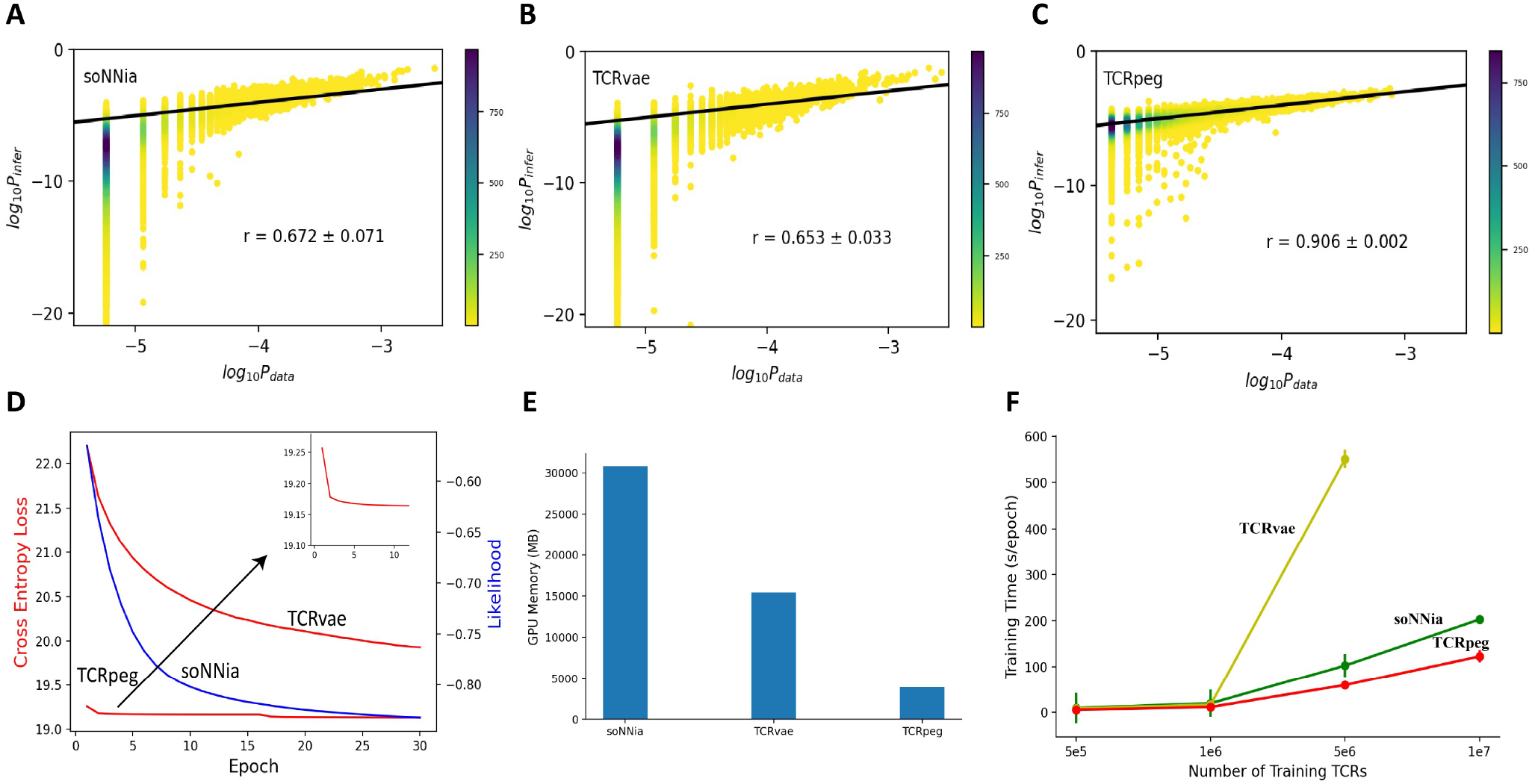
Performance of TCRpeg compared to the other two baseline methods soNNia and TCRvae. (**A-C**) Scatterplots of observed frequency *P_data_* vs. estimated probability *P_in f er_* for (**A**) soNNia, (**B**) TCRvae and (**C**) TCRpeg models trained on the large TCR pool combining 743 individuals from Emerson *et al*.^39^, along with the corresponding Pearson correlation coefficient *r*. The color indicates the number of sequences. (**D-F**) Comparison of soNNia, TCRvae, and TCRpeg model from practical aspects. Experiments are conducted under the same settings (learning rate and batch size) on a single Nvidia Tesla V100 GPU card with maximum 32 gigabytes memory. (**D**) The training curves for these three models. The soNNia model uses the likelihood as the model objective function (shown in the blue curve), while TCRvae and TCRpeg model minimize the cross-entropy loss (shown in the red curves). TCRpeg only needs less than ten epochs to converge, while the other two take around 30 epochs to converge. (**E**) The bar plot shows the GPU memory required to train each model. TCRpeg is more hardware-friendly. (**F**) The training speed of each model. TCRpeg takes less time to complete one training epoch compared to soNNia and TCRvae.

### TCRpeg helps profile TCR repertoires

The learned probability distribution can help profile the TCR subrepertoires in a probabilistic manner. Here, we were interested in learning the cell-type-level discrepancy and exploring the tissue-level differences since T-cells migrate and reside in different tissues and are influenced by different tissue environments. During maturation in the thymus, T-cells are selected and differentiate into two major cell types: cytotoxic (CD8^+^) and helper (CD4^+^) T-cells which function differently. Thus, our aim was to explore the TCR preferences of different TCR subrepertoires. To collect the data, we pooled TCRs with unique nucleotide sequences from nine healthy individuals from Seay *et al*.^21^. These TCR sequences were classified into three cell types (CD4^+^ conventional T-cells [Tconvs], CD4^+^ regulatory T-cells [Tregs], and CD8^+^ T-cells) and collected from three tissues (pancreatic draining lymph nodes [pLNs], mesenteric or inguinal “irrelevant” lymph nodes [iLNs], and spleen); that is, we have nine classes of subrepertoires. We applied TCRpeg to infer the probability distribution of each subrepertoire and quantified the difference between these distributions using the Jensen-Shannon divergence *D_JS_* (Methods).

We observed that the subrepertoires belonging to the same cell type are more conserved across different tissues. The same cell type in different tissues shows a lower TCR subrepertoire divergence, with an average Jensen-Shannon divergence as *D_JS_* ≃ 0.014 bits (Fig. 3A). However, the divergence is high between CD8^+^ and CD4^+^ TCR subrepertoires with the average *D_JS_* ≃ 0.041 bits. Tconv and Treg within the class of CD4^+^ cells demonstrate moderate similarities, with average *D_JS_* ≃ 0.024 bits. These observations confirm the results from Isacchini *et al*.^19^, where larger divergence between the CD8^+^ and CD4^+^ TCR subrepertoires and lower difference between the Tconv and Treg TCR subrepertoires are shown. These results were as expected since the CD8^+^ and CD4^+^ T-cells function significantly different: CD4^+^ T-cells are MHC-II restricted and pre-programmed for helper functions, whereas CD8^+^ T-cells are MHC I-restricted and pre-programmed for cytotoxic functions^40^. Additionally, subrepertoires of different tissues showed minor divergence, indicating that subsets of T cells perform similar functions across tissues.

**Figure 3.**
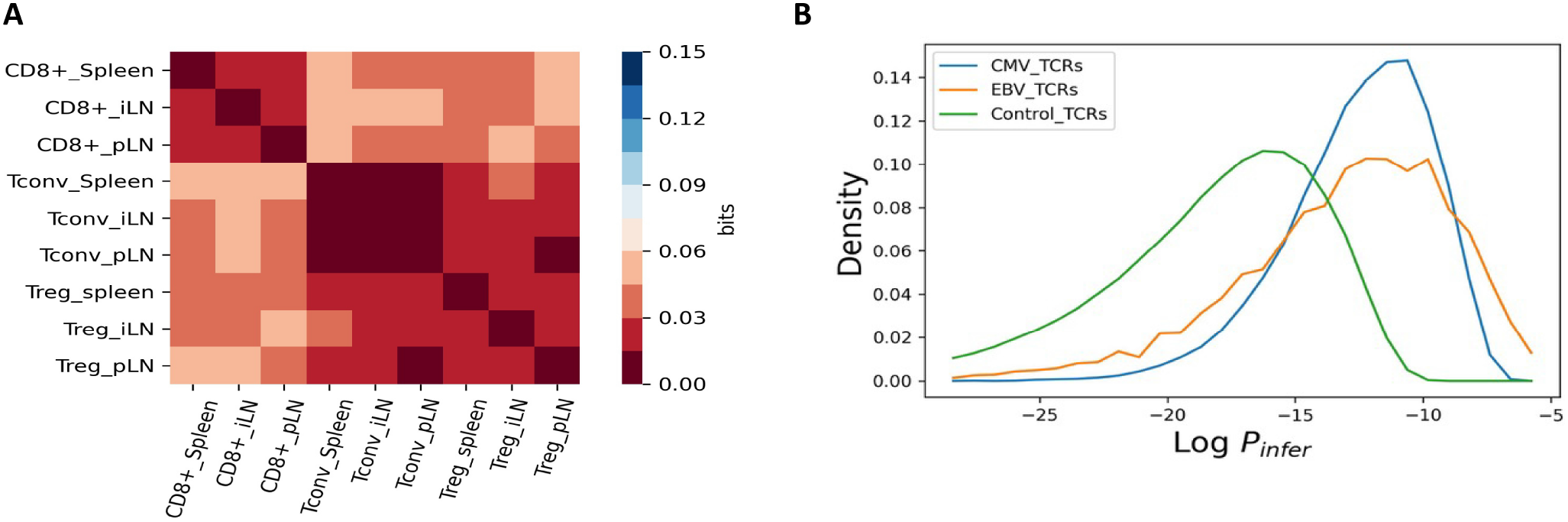
(**A**) Jesen-Shannon divergences between TCR subrepertoires at the cell type and tissue level. Jensen-Shannon divergences (*D_JS_*) were computed from TCRpeg trained on different subrepertoires (Methods). (**B**) Density map of inferred logarithmic probabilities for the three repertoires according to the TCRpeg model. For each repertoire, we inferred a TCRpeg model.

Next, we profiled two infection-specific TCR repertoires to further validate TCRpeg. We collected TCRs associated with cytomegalovirus (CMV) and Epstein-Barr virus (EBV) from VDJdb^41^ with 18,560 and 4,350 sequences, respectively. Furthermore, we randomly sampled 10^6^ TCRs from the aforementioned universal TCR pool as control. Figure 3B illustrates the density map of inferred probabilities for each repertoire. As expected, each repertoire had a distinct probability distribution. We then used a simple classifier to further show the characterization capacity of TCRpeg. We first trained a TCRpeg model for each repertoire. Then, we assigned a TCR *x* to the group *r* if *P_r_*(***x***) > *P_r′_* (***x***), and vice versa, where *r* and *r*′ are two repertoires. Interestingly, we observed an average accuracy 0.791 for classifying CMV-associated TCRs from control and 0.801 for classifying EBV-associated TCRs with a 5-fold cross-validation procedure.

### Classification of cancer-associated TCRs and SARS-CoV-2 epitope-specific TCRs

TCRpeg yields vector embeddings for TCRs sequences. Compared to the predefined or manually designed encoding method for TCR sequences, TCRpeg provides a learnable way to encode TCR sequences into vector representations. The update and reset gates of the GRU layers are learned during the training process to determine how much of the previous information stored in the hidden features needs to be passed along or abandoned^26^ (Fig. 1A). Therefore, the hidden features of the GRU layers at the last sequence position store summative information of the TCR sequences with different lengths; and these feature vectors provide an embedding for the TCR sequences.

To illustrate the embedding of the TCR sequence, we first collected cancer-associated TCR (caTCR) from Beshnova *et al*.^17^ (N~43,000) and SARS-CoV-2 epitope (YLQPRTFLL) specific TCRs from VDJdb database^41^ (N=683). We trained TCRpeg on these two datasets separately and obtained the respective numerical TCR embedding vectors. The UMAP dimensionality reduction^42^ was applied to project these vectors onto 2D space (Fig. 4A, Fig. 4B and Supplementary S3), showing that TCRs with a similar pattern (motif) tend to be clustered. It implies that the encodings could be helpful for antigen-specific TCR clustering. To further demonstrate the utility of TCRpeg-based encodings, we evaluated the classification performance on caTCRs and SARS-COV-2-epitope-specific TCRs using a fully connected neural network (FCN), taking these vector encodings as input. Since TCRpeg was designed mainly for TCR*β* chain and the paired TCRa and *β* chain are scarce, in this section we aimed mainly to investigate predictive performance with respect to TCR*β* sequences. We refer to this network as “TCRpeg-c” (Methods and Supplementary S4). To collect negative (or control) samples for the epitope-specific TCR dataset, we randomly sampled ten times more negative data than positive data from the universal repertoire of TCR mentioned above. We selected the CNN-based model, DeepCAT, developed in Beshnova *et al*. to compare with the caTCR prediction task, adopting the five-fold cross-validation procedure. In this prediction task, we observed an improvement in accuracy and predictive stability for TCRpeg-c with an average AUC ≃ 0.844 compared to DeepCAT with an average AUC ≃ 0.768 of caTCRs (Fig. 4C).

**Figure 4.**
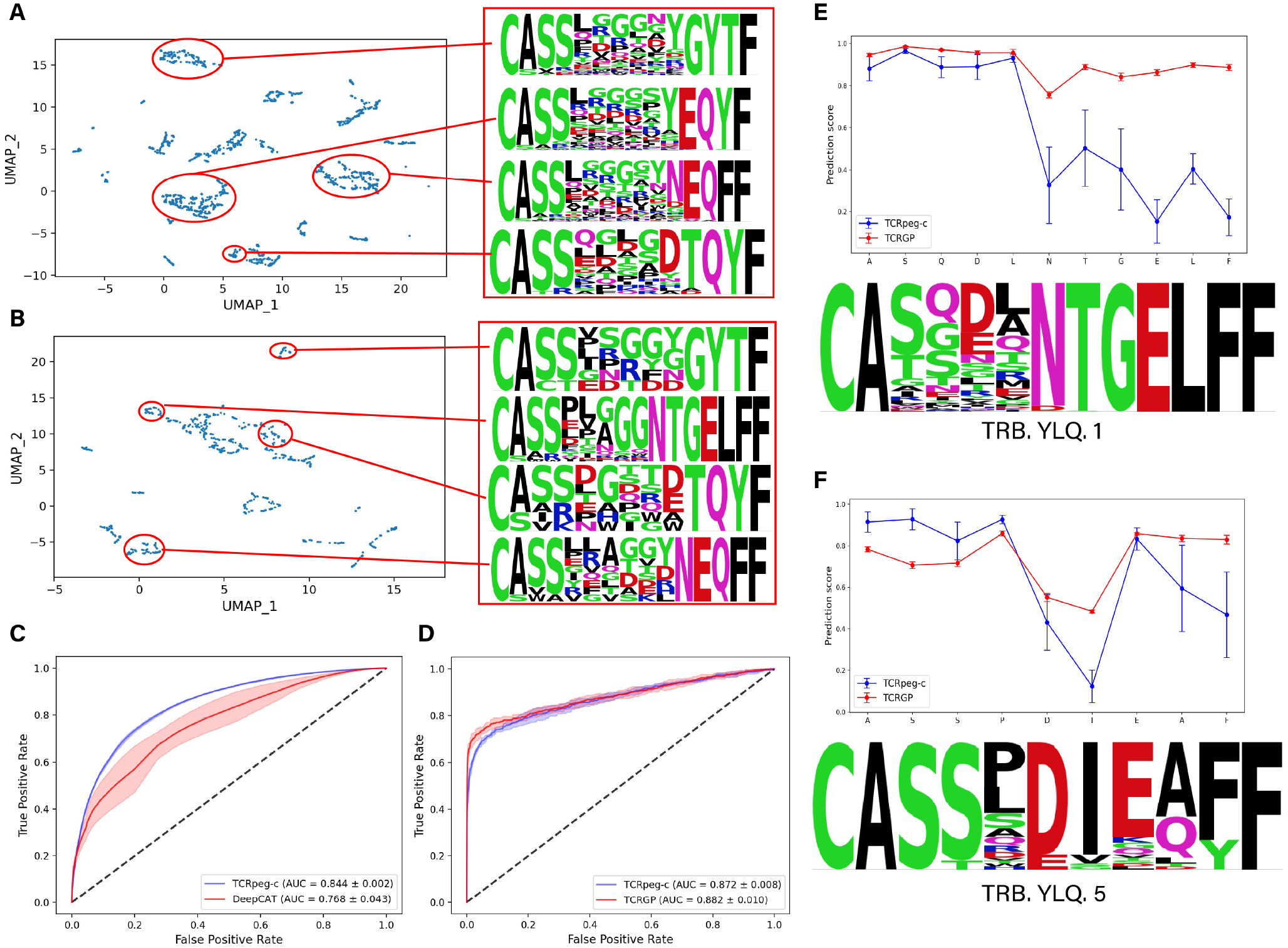
2D illustration of TCRpeg-based encodings and predictive performance for downstream classification tasks. (**A** and **B**) 2D projection map of encodings obtained from TCRpeg trained on (**A**) caTCRs and (**B**) specific TCRs of the SARS-CoV-2 epitope (YLQPRTFLL). More projecting results can be found in Supplementary S3. (**C** and **D**) ROC curves for tasks of (**C**) predicting caTCR and (**D**) SARS-CoV-2 epitope YLQPRTFLL. (**E** and **F**) Sensitivity analysis through amino acid substitutions used TCRpeg-c and TCRGP for two previously identified TCR motifs. For each position other than the two ends, we changed the amino acid at that position to the four other most frequent AAs and used these two models to score the modified sequences. TCRpeg-c is more sensitive than TCRGP to substitutions of amino acids inside the motifs.

In the more challenging epitope-specific TCR prediction task with scant data, TCRpeg-c still demonstrated competitive performance with AUC ≃ 0.872 compared to the baseline method TCRGP^27^ with AUC ≃ 0.882 (Fig. 4D). However, the TCRGP model is sophisticated, and it is designed specifically for the TCR-epitope mapping problem with low data size, combining multiple techniques including alignment of TCR sequences, Gaussian process (GP) and variational inference.

TCRpeg-c finds TCR motifs through perturbation analysis. TCR motifs are important and instructive in determining their specificity to antigens^43^. Previously, motif discovery for TCR repertoires was mainly accomplished by exploring similarities between TCRs such as the TCRNET method^44–46^ or investigation of frequency enhancement of k-mers for TCRs^43^. Here, we used predictive models to test the sensitivity of previously identified TCR motifs for specific TCRs of the SARS-CoV-2 epitope YLQPRTFLL (Methods). We observed the correspondence between previously identified TCR motifs and sensitive residues according to the TCRpeg-c predicted scores, indicating the importance of TCR motifs for epitope binding (Fig. 4E and 4F). However, although TCRGP achieves high predictive performance, it lacks the ability to detect sensitive residues (Fig. 4E and 4F). We attribute its insensitivity to the need to pad TCR sequences to a fixed length, which could lower the degree of variation caused by amino acid substitution.

### Generating more TCR sequences with potentially the same specificity

A good generative model could be beneficial for the adoptive transfer of TCR engineered T-cells (TCR-T) that has been applied to treat viral infections such as hepatitis B and C^47, 48^, cancer immunotherapy^49, 50^, and autoimmune disease therapy^51^ through *in silico* generation of similar TCR sequences guiding the in vitro TCR design. We extended TCRpeg to be generative through a simple sampling strategy (Methods).

We first aimed to systematically evaluate and compare the generation ability of TCRpeg with the baseline methods, soNNia and TCRvae, in terms of the statistical properties between the generated TCR sequences and real sequences. Specifically, we investigated the distributions of sequence lengths, positions of amino acids, V gene and J gene usages. We observed strong agreement between the probability distributions of *in silico* and real TCR repertoires for both the TCRpeg and soNNia models. For the position distributions of each amino acid, the TCRpeg- and soNNia-generated sequences successfully fitted the original statistics with an average Pearson correlation coefficient *r* ≃ 1.0 and *r* ≃ 0.999, respectively, compared to TCRvae with *r* ≃ 0.982 (Supplementary S6). For V and J gene usages, the TCRpeg and soNNia models still outperform TCRvae, achieving average *r* ≃ 0.999 and *r* ≃ 0.998 compared to *r* ≃ 0.949 of the TCRvae model for V gene usage distribution (Fig. 5A and Supplementary S7), and *r* ≃ 1.0 and *r* ≃ 0.997 over *r* ≃ 0.993 for J gene usage distribution (Fig. 5B). For the length distribution, these three models all achieved highly accurate performance with r ≃ 1.0,0.998,1.0 for TCRpeg, soNNia, and TCRvae, respectively (Fig. 5C). The generation performance of TCRpeg is stable and accurate even when trained on a small subset of TCRs (Supplementary S8 and S9). These results together highlight that TCRpeg is reliable for summarizing a TCR repertoire, and consequently, generating new sequences in recovering the real statistical distributions.

**Figure 5.**
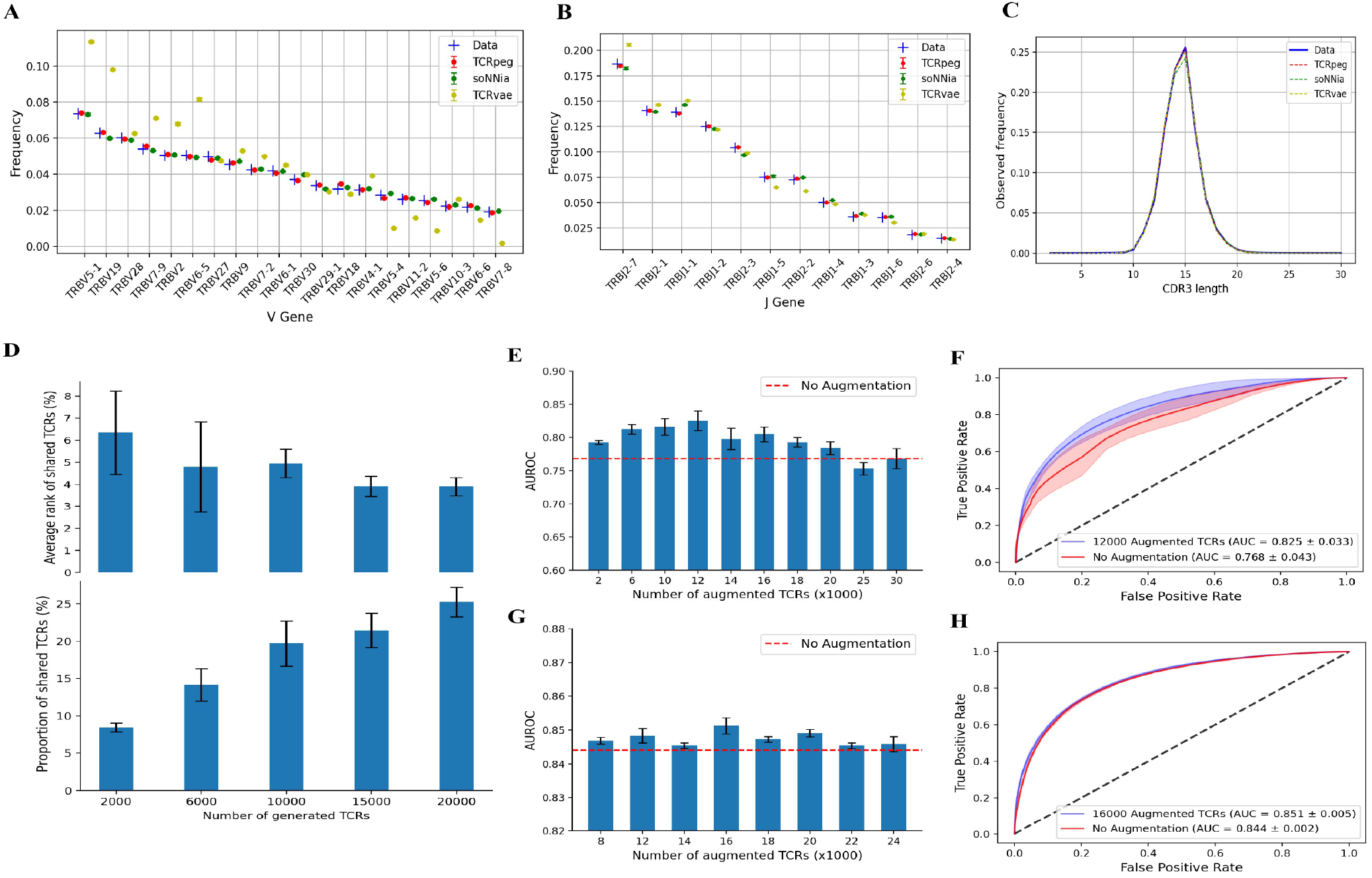
Characteristics of the TCR sequences generated by the three generative models. (**A**-**C**) Comparison of the statistical distributions of the generated sequences with the real data with respect to (**A**) V gene usage, (**B**) J gene usage and (**C**) length distribution. In (**A**), only the top 20 frequent V genes are listed. We include the figure of full V gene usage and the distributions of amino acids in Supplementary S7 and S6. (**D**) The proportion of the TCR sequences in the test set that also appears in the generated TCRs (bottom panel) and the average probability rank of those shared TCRs among the generated TCRs (top panel). With more TCRs being generated, more of them can be found in the test set. (**E** - **H**) Performance in the task of predicting caTCRs by applying the TCR-specific data augmentation technique. (**D** and **F**) The AUC scores with a different number of augmented TCR sequences when using the (**E**) DeepCAT model and (**G**) TCRpeg-c. (**F** and **H**) ROC curves for the DeepCAT model (**E**) and TCRpeg-c **G** with the best number of augmented TCRs.

A reliable generative model should be able to produce new TCR sequences with “hidden similarity” to real TCR data, in addition to statistical similarity. Here, we were interested to determine whether the generated TCR sequences possess the same epitope specificity with the data used in training TCRpeg. To verify this, we retrained TCRpeg on the training set of the TCRs specific to the epitope YLQPRTFLL and utilized it to generate new sequences accordingly. We first noticed that some of the TCRs in the test set could also be found in the generated data set (Fig. 5D), which shows the generative power of TCRpeg given the wide potential diversity of TCR sequences. To take a closer look at these generated TCR sequences, we observed that those TCRs that were also found in test set possessed high generation probabilities (averagely ranked < 10% among generated sequences, Fig. 5D). Finally, we utilized the TCRMatch^28^ software to further validate the hidden similarity of the generated TCR sequences and observed that 50 – 60% of them possess the same epitope specificity as the TCR sequences used in training according to a scoring threshold of 0.9 (Supplementary S10). On the contrary, although the soNNia and TCRvae models achieve comparable performance with respect to statistical similarities, only less than 40% of the generated sequences possess the same epitope specificity determined by the same scoring threshold (Supplementary S10). Overall, our results indicate that TCRpeg can generate new TCR sequences with statistical and possible hidden similarities to the TCRs used for training.

### Augmenting TCR sequencing data

Data augmentation techniques are ubiquitously used in machine learning tasks to increase the generality of data by adding similar samples generated by either slightly modifying the original data or synthesizing similar data. They act as regularizers to alleviate the issue of overfitting and improve the generalization capacity of machine learning models, especially when applied to computer vision tasks^52^ or natural language processing tasks^53^. Adopting the data augmentation techniques here should improve the classification of TCR sequences. TCR sequences might abolish their epitope specificity by amino acid substitutions, especially when they happen inside contact motifs^43, 54^; therefore, directly performing amino acid substitutions, insertions, or deletions on TCR sequences cannot work as data augmentation. However, with strong generative ability, TCRpeg may generate similar TCR sequences and serve as a computational tool for TCR-specific data augmentation.

To analyze the feasibility of TCR-specific data augmentation, we evaluated and compared the predictive performance of classifying caTCRs with and without data augmentation while keeping all other training settings unchanged. For the DeepCAT model, we observe a large performance gain with up to 0.057 higher AUC when applying data augmentation technique (Fig. 5E and 5F). For the TCRpeg-c model, we still find accuracy enhancement in the AUC value from 0.844 to 0.851 with data augmentation (Fig. 5G and 5H). Besides, the AUPRC (area under the precision-recall curve) also increases and the test loss decreases, which is a positive sign of mitigation of overfitting (Supplementary S11). To further validate the utility of our TCRpeg-based augmentation technique, we performed classification for the influenza epitope GILGFVFTL and EBV epitope GLCTLVAML specific TCRs with 3,406 and 962 positive samples, respectively, using the TCRex model^55^. Without changing any training settings, we observed up to 2.1% and 21.4% accuracy enhancement for these two TCR datasets (Supplementary S12).

## Discussion

An accurate probabilistic model for large-scale TCR sequencing data is a cornerstone for a better understanding of functional TCR repertoire. Previous works have developed selection models soNia^25^, soNNia^19^, and the VAE-based model TCRvae^29^ to characterize the distribution of productive TCR sequences. However, they are all intrinsically unable to capture the information behind the length variation. In this work, we introduced TCRpeg, an autoregressive deep learning model that utilizes a recurrent neural network with GRU layers to characterize the TCR repertoires. Unlike soNia, soNNia, and TCRvae which need to pad every TCR sequence to the same length, TCRpeg can process TCR sequences with any lengths. Such capability can eliminate the noise introduced by adding an extra “amino acid” for padding and take advantage of the information behind the variance in lengths.

We first demonstrated that TCRpeg can improve the statistical characterization of TCR repertoires in a large cohort of individuals^39^ compared to soNNia and TCRvae by a large margin, which implies that TCRpeg can better learn the TCR sequence pattern. We attribute the superior performance of TCRpeg to its ability to process TCRs with different lengths and its transmission of hidden features that properly store the previous information. In particular, TCRpeg takes less iterations to converge and requires lower computation resources. These results indicate the advantages of using an autoregressive model that is capable of processing TCR sequences with different lengths to describe large-scale TCR sequencing data from a probabilistic perspective.

Using the statistical inference power of TCRpeg, we explored the differences and similarities between functional TCR subrepertoires collected from different T-cell types or tissues at the repertoire level. We discovered that TCR subrepertoires belonging to families with more closely related developmental paths (i.e., Tconvs and Tregs) possess higher statistical similarities. Meanwhile, they both show large differences with CD8^+^ T-cells that diverged earlier in T-cell maturation. Next, we explored the statistical profile of the infection-specific TCR repertoires and observed their distinct patterns through the density map (Fig. 3B). To illustrate the characterization capacity of TCRpeg in a more straightforward way, we used a simple classifier that directly applied the probability inference ability of TCRpeg to classify the infection-specific TCRs. This simple classifier achieved relatively high prediction performance, with an average accuracy of 0.791 for classifying CMV and 0.801 for classifying EBV associated TCRs. Our results showed that TCRpeg is a superior tool for characterizing TCR repertoires from a statistical perspective.

On the basis of the architecture of TCRpeg, we can obtain helpful vector representations of TCR sequences from the trained TCRpeg model, which is not provided by soNNia or TCRvae. Compared to other predefined or hand-designed encoding methods for TCR sequences, TCRpeg provides a learnable way to encode TCR sequences by updating functional gates inside GRU layers^26^. We observed that TCRpeg-based TCR encodings could reflect the degrees of similarities between TCR sequences that sequences with a similar pattern (motifs) tend to cluster together (Fig. 4A and 4B). This suggests a potential application of antigen-specific TCR clustering, since shared TCR motifs indicate the same antigen specificity.

To examine the performance of TCRpeg-based encodings in a predictive manner, we assessed the classification performance of caTCRs and YLQPRTFLL epitope-specific TCRs using a fully connected neural network taking these vector encodings as input (TCRpeg-c). For the caTCR prediction task, we chose the DeepCAT model developed by Beshnova *et al*. as the baseline method. We observed a significant improvement in accuracy and predictive stability for TCRpeg-c compared to DeepCAT in the prediction of caTCRs (Fig. 4C). With such high precision, TCRpeg-c could facilitate cancer detection through the process introduced in Beshnova *et al*.. In recent years, multiple machine learning methods have been developed to predict the epitope specificity of TCRs, such as TCRex^55^, DeepTCR^54^, and TCRGP^27^. All of these methods have explored the problem in slightly different settings and compared with each other. In the more challenging classification task of predicting SARS-SoV-2 epitope (YLQPRTFLL)-specific TCRs, we compared TCRpeg-c to a representative of the above group of machine learning models, TCRGP, which is a combination of multiple functional modules including TCR alignment, Gaussian process, and variational inference. TCRpeg-c demonstrated competitive performance in this task compared to TCRGP (Fig. 4D). In particular, TCRpeg-c is sensitive to substituting for an amino acid primarily when it occurs inside the TCR motifs, while TCRGP is insensitive to that (Fig. 4E and 5F). This finding indicates that TCRpeg-c can be used for motif validation and help with TCR engineering for immunotherapies^56^. In addition, this perturbation analysis might reveal *de novo* motifs that have not yet been discovered using nonpredictive methods (Supplementary S5). The comparable accuracy performances in the above two classification challenges validate the advantage of TCRpeg-based encodings, which can be further concatenated with epitope features to facilitate the unseen epitope-TCR interaction prediction task^57^.

One direct application of TCRpeg is to generate new TCR sequences with characteristics similar to those of natural sequences. We first compared the generation capability of TCRpeg with soNNia and TCRvae with respect to the statistical distributions on the large universal TCR pool we have constructed. We showed that TCRpeg-generated TCR sequences had the closest amino acid distributions, length distribution, and V/J gene usages to the real sequences compared to the other baseline methods. Next, we found that some TCRs in the test set could also be found in the generated dataset, and those shared TCRs have high generation probabilities among the generated dataset (Fig. 5D). These results imply that newly generated TCR sequences with high probabilities might share the same epitope specificity with the data used in training, providing a potential way to meet the demand for more data. We further applied the TCRMatch^28^ software to validate this implication and show that 50 – 60% of the generated TCRs share the same epitope specificity as the TCRs used for training. On the contrary, less than 40% of the TCRs generated using TCRvae or soNNia share the same specificity (Supplementary S10). The generative power of TCRpeg can also be used to design similar TCRs to facilitate immunotherapy for T-cell transfer^49–51^.

Data augmentation is a ubiquitous technique used to increase the performance of machine learning models, especially in computer vision systems^52^. Given that more and more machine learning models have been developed for TCR-related tasks and the acquisition of more data is costly and time consuming, which restricts the development of highly accurate machine learning models, we developed and validated the TCR-specific data augmentation technique empowered by TCRpeg to relieve such restriction. For the caTCR classification task, we observed a notable improvement with data augmentation (Fig. 5E and 5F). In addition, we further validated the utility of data augmentation using another machine learning model - TCRex^55^ in the prediction tasks of GILGFVFTL and GLCTLVAML specific TCRs and again observed an improvement in accuracy (Supplementary S12). However, in the SARS-CoV-2 specific TCR recognition task, data augmentation failed to boost the model performance. When learning from such a small data size, TCRpeg tends to generate highly similar TCRs with those in the training set and thus provides limited additional information to the predictive model, which might result in more severe overfitting. Nevertheless, TCRpeg-based data augmentation is a free option for boosting model performance without any extra cost.

In this work, we have introduced a new holistic software tool TCRpeg for estimating the probability distribution of a TCR repertoire with a great performance enhancement over previous works. Furthermore, with promising performance in probability inference, TCRpeg improves on a range of TCR-related tasks: (i) reveal TCR repertoire-level discrepancies from a probabilistic prospective; (ii) classify antigen-specific TCRs and validate previously discovered TCR binding motifs; (iii) generate novel TCRs and augment TCR data for accuracy enhancement of machine learning models. Our results and analysis highlight the flexibility and capacity of TCRpeg to extract TCR sequence information, providing new insights for understanding the complex genomic concepts hidden behind TCR repertoires.

## Methods

### Data Description

The data sets used in this work are classified into three groups to evaluate the performance of TCRpeg. We filter out TCRs with lengths greater than 30 or not starting with a cysteine in all data sets. We also verified sequences that are written as V gene, CDR3 sequence, J gene and removed sequences with unknown genes. In addition, we only considered the 20 standard amino acids in this work and removed sequences with any unspecified amino acid. The detailed descriptions of each group of data are shown below:

1. To quantify the precision of the inference of TCRpeg along with the other two baseline methods, we used the TCR repertoires sampled from a large cohort, including 743 individuals from Emerson *et al*.^39^ We pooled the unique nucleotide sequences of receptors from all individuals and built a universal TCR pool that contains around 10^9^ sequences in total. The multiplicity of an amino acid sequence in this universal TCR pool indicates the number of independent recombination events that led to that receptor. We randomly and equally split the TCR pool into a training set and a test set.
2. To characterize the differences between the TCR subrepertoires of functional cell types collected from different tissues, we pooled unique TCRs from 9 control donors from Seay *et al*.^21^ at the tissue level. These TCR sequences were sorted into three cell types and collected from three tissues. Thus, for each donor status (healthy or T1D), we have nine groups of TCRs. Again, the multiplicity of an amino acid sequence in this universal TCR pool indicates the number of independent recombination events that led to that receptor, which is used to calculate the real probability distribution.
3. To evaluate the performance of TCRpeg-c in classification tasks, we first collected cancer-associated TCRs (caTCRs) from Beshnova *et al*.^17^. Briefly, Beshnova and his colleagues collected TCR sequences from approximately 4,200 recorded samples downloaded from The Cancer Genome Atlas (TCGA) and excluded those sequences that are also found in healthy donors. The remaining around 43000 TCR sequences are assumed to be cancer-associated TCRs (caTCRs). We extracted the SARS-CoV-2 epitope (YLQPRTFLL), influenza epitope GILGFVFTL and EBV GLCTLVAML specific TCRs from VDJdb^41^ database (positive TCRs N = 683, 3406, 962, respectively, extracted on 24 January 2022). We then randomly sampled ten times more negative data than positive data from the universal TCR pool constructed previously to serve as the control TCRs.

### TCRpeg and TCRpeg-c

The illustrations of TCRpeg and TCRpeg-c are shown in Fig. 1A and Supplementary S4. To enable the training of TCRpeg, we first trained the word2vec^38^ model on 1 × 10^6^ TCR sequences randomly sampled from the pooled universal repertoire aforementioned to obtain the numerical embeddings for each amino acid, regarding the amino acid as the “words” and the TCR sequences as the “sentences”. Specifically, we adopted the skip-gram architecture with the window size and embedding size set to 2 and 32 and trained it for 20 epochs. For the TCRpeg model, the GRU modules have three layers with the size of the hidden feature set to 64. We trained TCRpeg using the Adam^58^ optimizer for 20 epochs to minimize the cross-entropy loss between the soft-maxed logits and the one-hot encoded representation of the discrete categorical outputs of the network. The probability of a given TCR sequence *P_in f er_*(***x***) is estimated using Equation 1. Specifically, we input the given TCR sequence to TCRpeg and obtain the corresponding output probability distribution of the amino acid at the next time step. Thus, *P_in f er_*(***x***) is the multiplication of the probabilities of amino acids at each time step.

For the TCRpeg-c model, the size of the hidden feature is increased to 512 to better capture the hidden sequence features for classification tasks. On top of the pre-trained TCRpeg, the fully connected neural network contains two hidden layers with 384 and 96 neurons, followed by the ReLU activation function. In the task of predicting caTCRs, we trained TCRpeg-c for 30 epochs to minimize the loss of cross-entropy between the output logits and true labels, with dropout operations (p=0.2) to reduce the issue of overfitting. In classifying epitope-specific SARS-CoV-2 TCRs, we trained TCRpeg-c for 20 epochs with a dropout rate set to 0.4. In both above-mentioned classification tasks, the TCRpeg was trained on the respective training set to provide the numerical embeddings for TCRs. The trained TCRpeg-c can be used to find TCR motifs through perturbation analysis. Specifically, we permuted each position of the TCR sequences except for the first and last positions, with four other amino acids that most likely appeared at that position according to the amino acid frequency at that position. We adopted this strategy to avoid skewed permuted sequences containing amino acids at some positions with nearly zero probabilities. We then applied the trained TCRpeg-c to score each permuted sequence to determine residues that are sensitive to changes.

### Quantifying the accuracy of probability inference

To evaluate the precision of probability inference, we compared the estimated probabilities *P_in f er_*(***x***) to the observed frequencies *P_data_*(***x***) of the test set. The accuracy can be quantified by Pearson’s correlation coefficient *r* between *P_in f er_*(***x***) and *P_data_*(***x***). A higher value of *r* indicates a better model. The calculation of *P_in f er_*(***x***) for TCRpeg is described in the previous section using the autoregressive likelihood formula. For the two baseline methods TCRvae and soNNia, we compute *P_in f er_*(***x***) by:

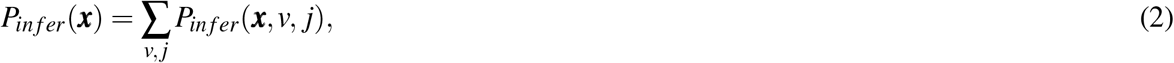

which sums the V and J genes along with the TCR sequence ***x***. Finally, we normalize the inferred probabilities *P_in f er_*(***x***) and consider them as the approximation of the real probability distribution.

### Quantifying of difference between TCR subrepertoires

We used the Jensen-Shannon divergence *D_JS_*(*r^i^, r^j^*) to characterize the difference between two TCR subrepertoires *r^i^* and *r^j^*:

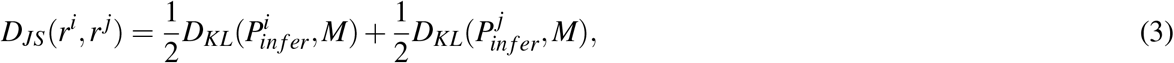

where 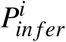 and 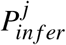 are computed by two different TCRpeg separately trained on subrepertoires *r^i^* and 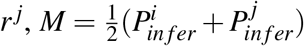 and *D_KL_* represent the Kullback-Leibler divergence. To characterize the differences between the TCR subrepertoires of functional cell types collected from different tissues, we first trained TCRpeg on each tissue-level TCR subset for 20 epochs with hidden size and the number of layers set to 128 and 3, respectively. Then we applied Eq. 3 to calculate the JS divergences between each pair of those TCR subrepertoires.

### Using TCRpeg to generate TCR sequences

We adopted a simple sampling method to generate new TCR sequences using TCRpeg. Specifically, we first input the start token (“<SOS>”) to the TCRpeg and then randomly sampled the amino acid for the next position from the output probability distribution (computed using the Softmax operation). Following the same procedure, at each time step, we randomly sampled the amino acid for that time step according to the probability distribution defined by the predicted scores and input it to the next time step to obtain the following amino acids. This stochastic generation procedure can be described by the formula stated below:

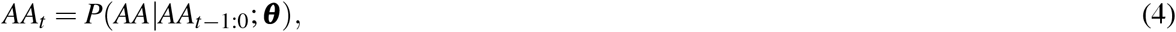

where *AA*_0_ stands for the start token and ***θ*** represents the TCRpeg parameters. The generation process stops when the special stop token (“<EOS>”) is generated. To allow the ability to infer the corresponding V and J gene along with the TCR sequence, we extended TCRpeg and formulated the probability of a given TCR sequence ***x*** with specific V and J genes as:

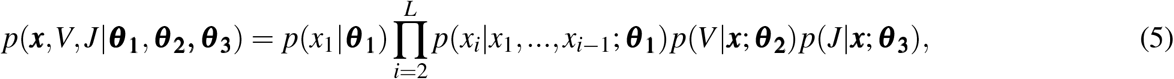

where *p*(*V*|***x***; ***θ*_2_**) and *p*(*J*|***x***; ***θ*_3_**) are the probabilities conditioning on the TCR sequence ***x***; ***θ*_2_** and ***θ*_3_** are parameterized by two respective fully connected single-layer neural networks. The TCRpeg, soNNia and TCRvae models were inferred from the universal TCR repertoire aforementioned, and then we applied them to generate new TCR sequences along with V and J genes.

## Availability of data and materials

All data analyzed in this work can be found in the original publications that collected the data^17, 21, 39, 41^, and we include the preprocessed data at https://github.com/jiangdada1221/TCRpeg#data. TCRpeg was written in Python using the deep learning library Pytorch^59^ and is available as a python package. Source code, use-case tutorials, and documentations can be found at https://github.com/jiangdada1221/TCRpeg. Users can install directly from Github or PyPI via pip.

