## Supplementary Materials for "Deep autoregressive generative models capture the intrinsics embedded in T-cell receptor repertoires"

### 1 Supplementary

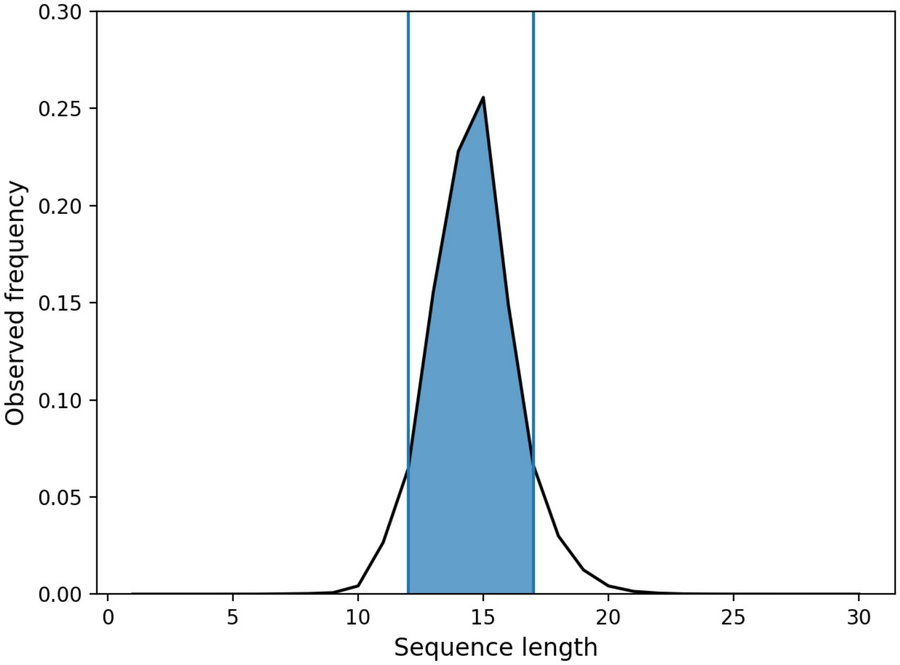


**Figure S1.** Length distribution of TCR sequences in our constructed universal TCR pool. The lengths of TCRs mainly range from 12 to 17, which take up 84% of the total TCRs (shown in blue shadow). This range is suitable for inferring a deep autoregressive model without causing the issue of gradient explosion or gradient vanishing.

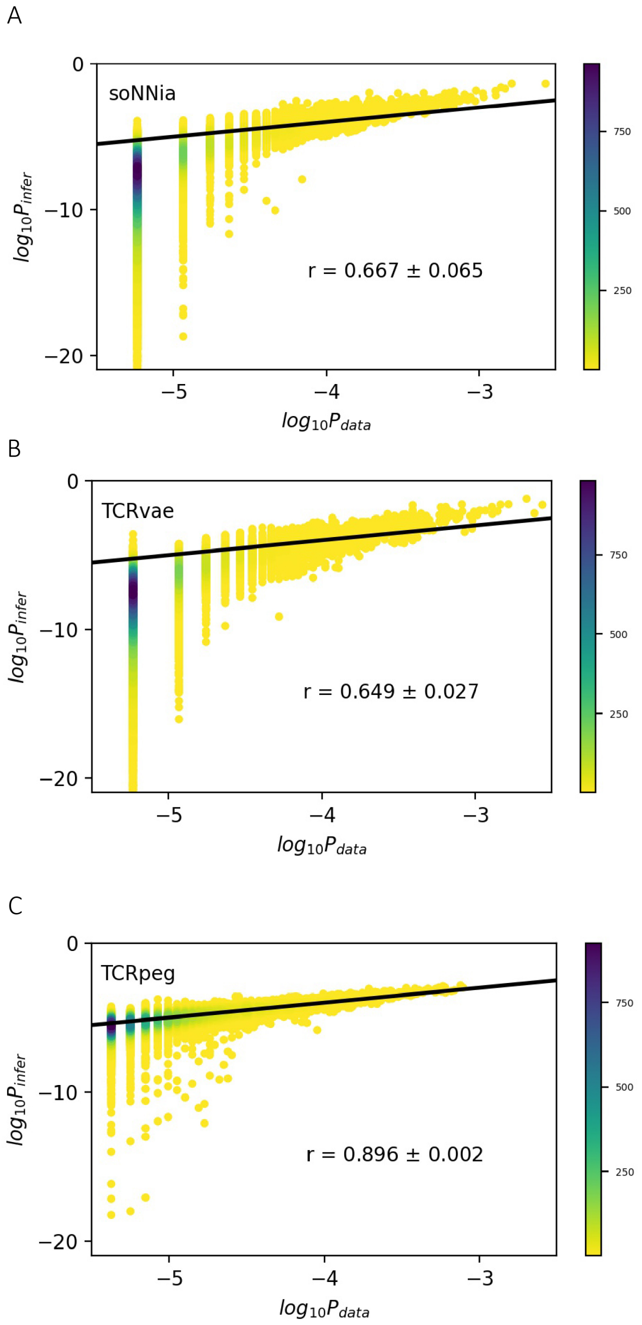


**Figure S2.** Inference performance of the soNNia, TCRvae, and TCRpeg models in a small proportion of the large universal TCR pool. To assess the stability of these generative models, we re-trained each of them on 200 thousand TCR sequences randomly sampled from the training set. We evaluated them on the test set under the same training settings. The results show that all these models are stable in probability inference, and TCRpeg still surpasses the other two baseline models by large margins.

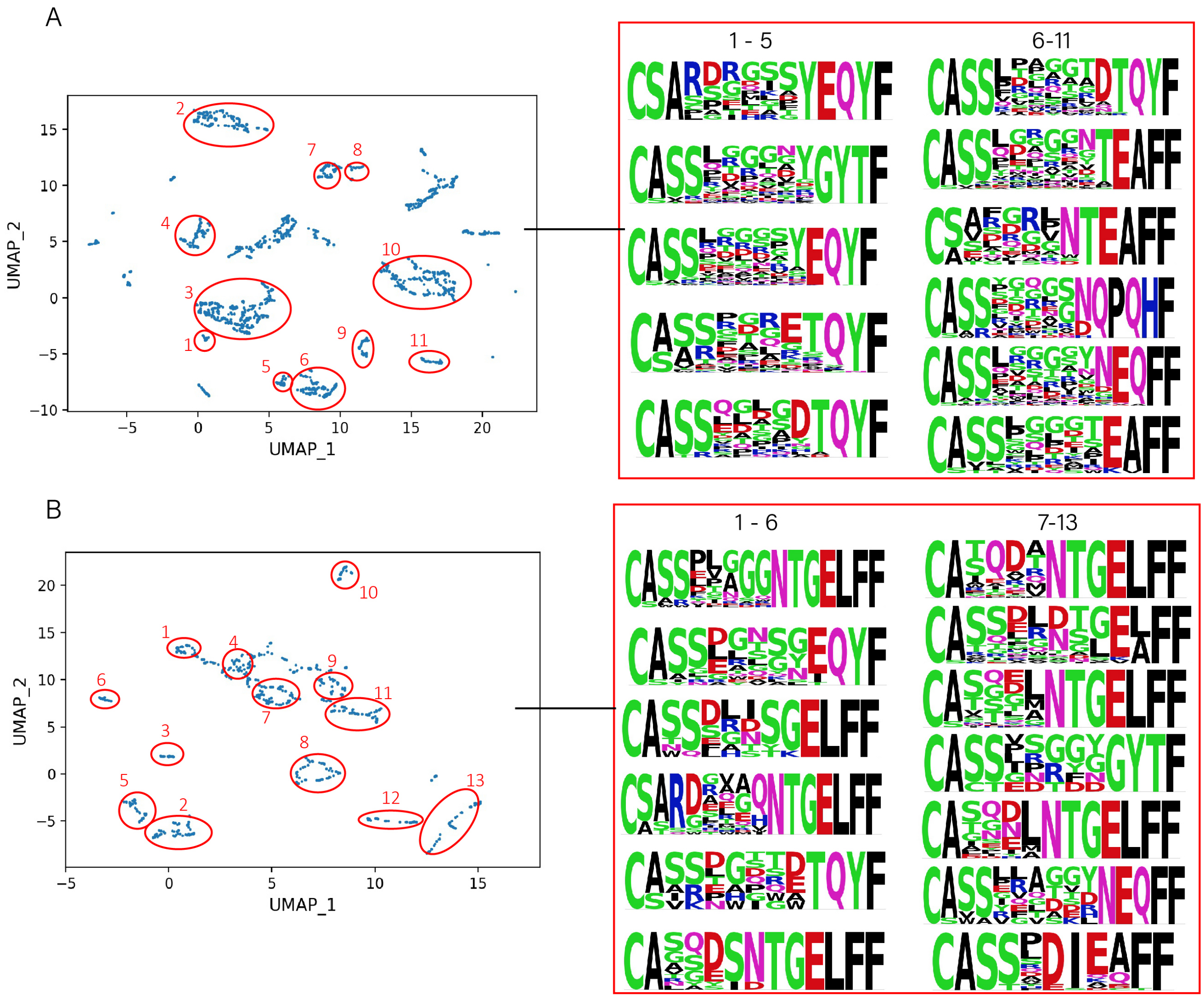


**Figure S3.** 2D projection map of TCRpeg-based encodings of (**A**) caTCRs and (**B**) YLQPRTFLL specific TCRs. We demonstrate more TCR patterns corresponding to different clusters in the projection map that are not shown in Fig. 4 due to the size limit.

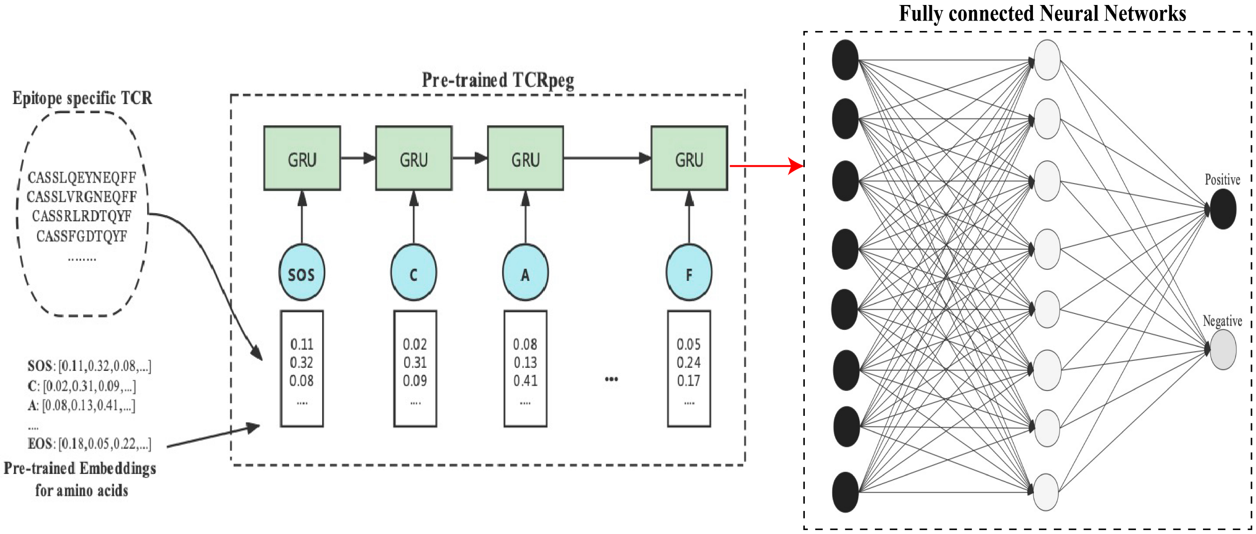


**Figure S4.** The illustration of the architecture of TCRpeg-c. First, we trained the TCRpeg on epitope-specific TCR sequences (or other specific TCRs) to obtain the numerical encodings for each TCR. Then these encodings were inputted into a fully connected neural network for classification purposes.

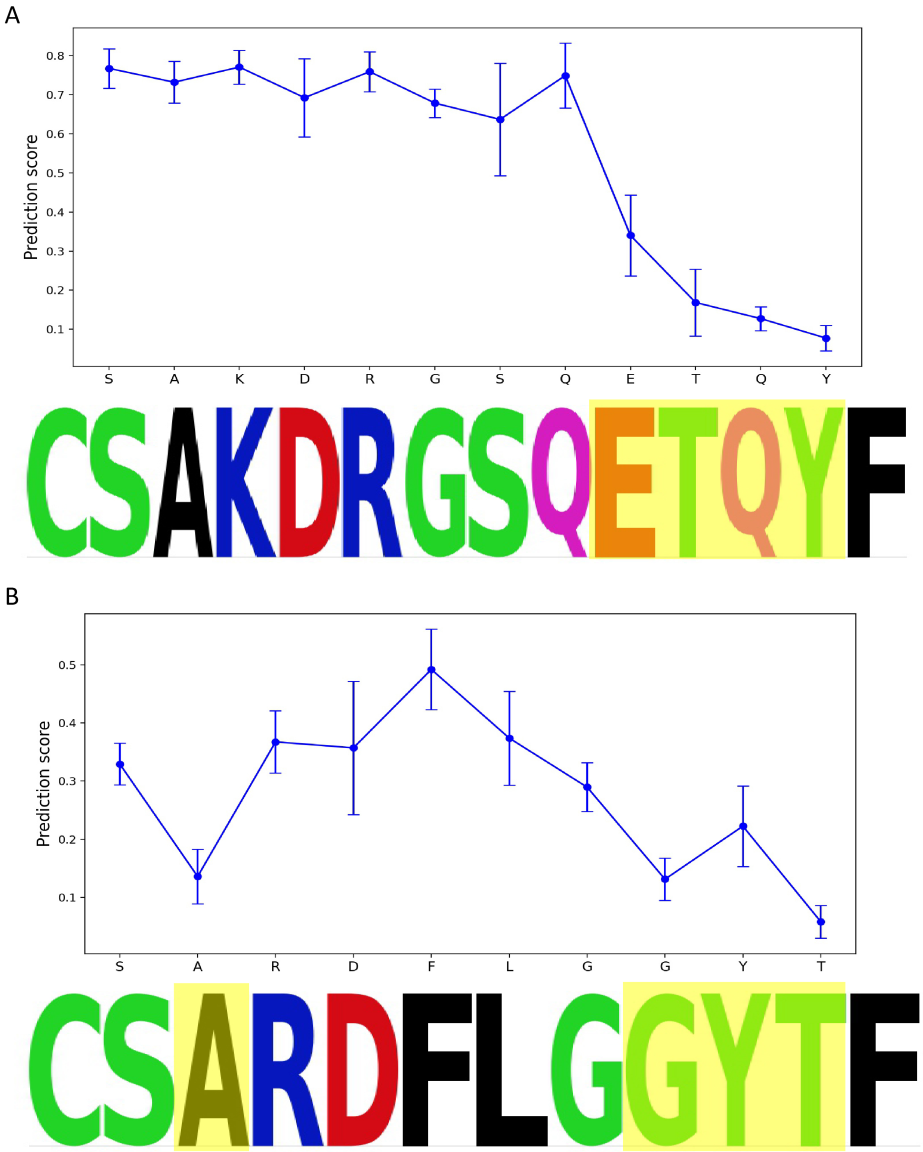


**Figure S5.** Perturbation analysis to discover *de novo* motifs. In addition to validating previously identified motifs, we performed a perturbation analysis on some TCRs to find potential motifs. Blue curves indicate the prediction scores from TCRpeg-c for the permuted TCR sequences. We observe dropoffs in the region of "ETQY" within the TCR sequence "CSAKDRGSQETQYF" (**A**) and "GYT" as well as "A" within "CSARDFLGGYTF" (**B**). We wish that when more YLQPRTFLL-specific TCRs are available, these two motifs could be identified through the TCR similarity network.

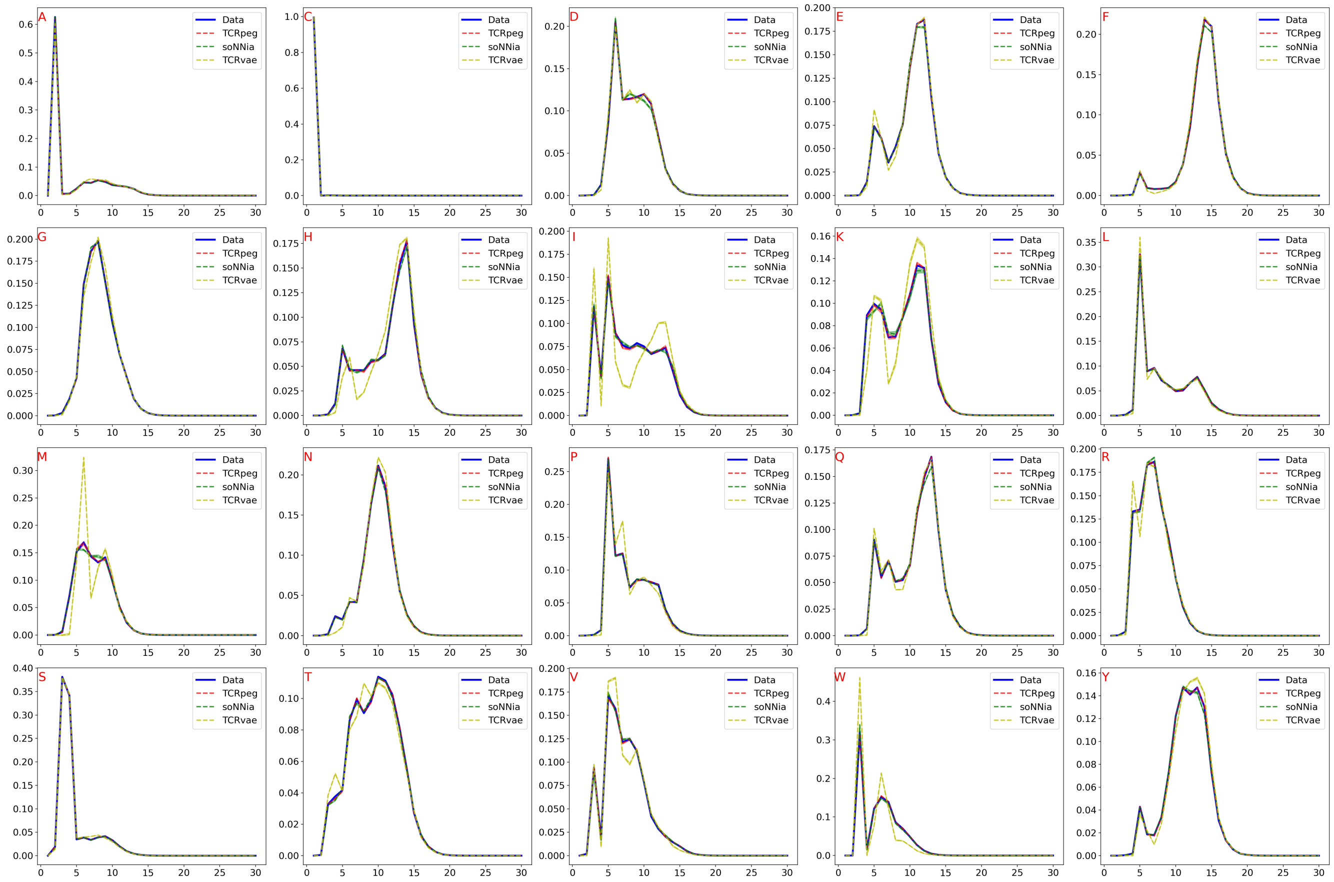


**Figure S6.** The position distributions of amino acids for the generated TCR sequences compared to TCRs from the universal TCR pool. Each subplot shows the position distribution of a particular amino acid. The x-axis represents the TCR sequence's position index (from left to right), and the y-axis shows the amino acid frequency appearing at each position index. The TCRvae model performs the worst in this task. The generated TCR sequences using the soNNia and TCRpeg models possess a close distribution to the observed data, and TCRpeg performs slightly better than soNNia.

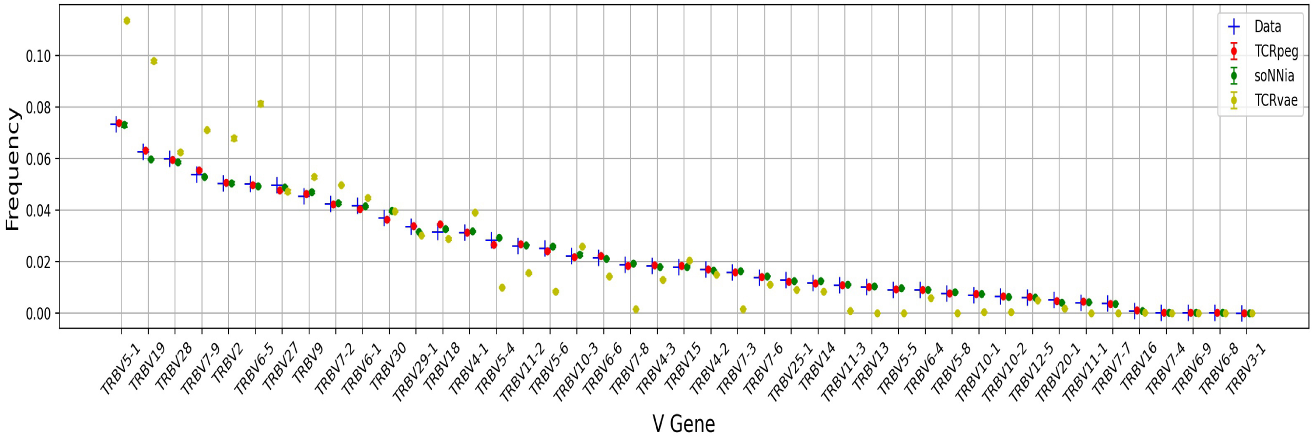


**Figure S7.** The full version of Fig. 5A, showing the statistical distributions of all 43 V genes from the generated repertoires using the three generative models.

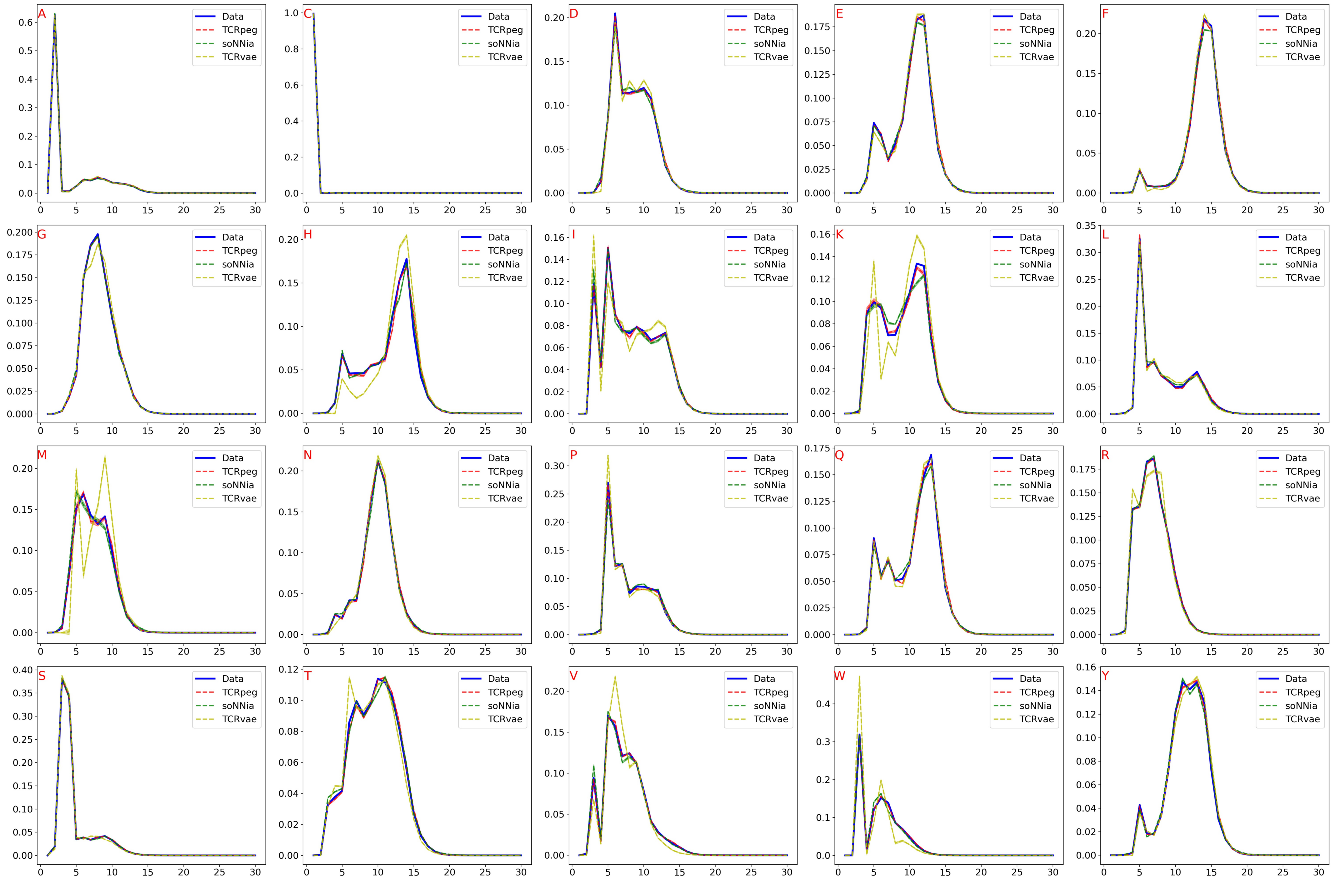

**Figure S8.** Position distributions of amino acids for the generated TCR sequences compared to TCRs from the universal TCR pool. We randomly sampled 200 thousand TCRs from the TCR pool and trained each model on this subset. TCRpeg still achieved the best performance with $r \simeq 0.999$ compared to $r \simeq 0.998$ for soNNia and $r \simeq 0.982$ for TCRvae.

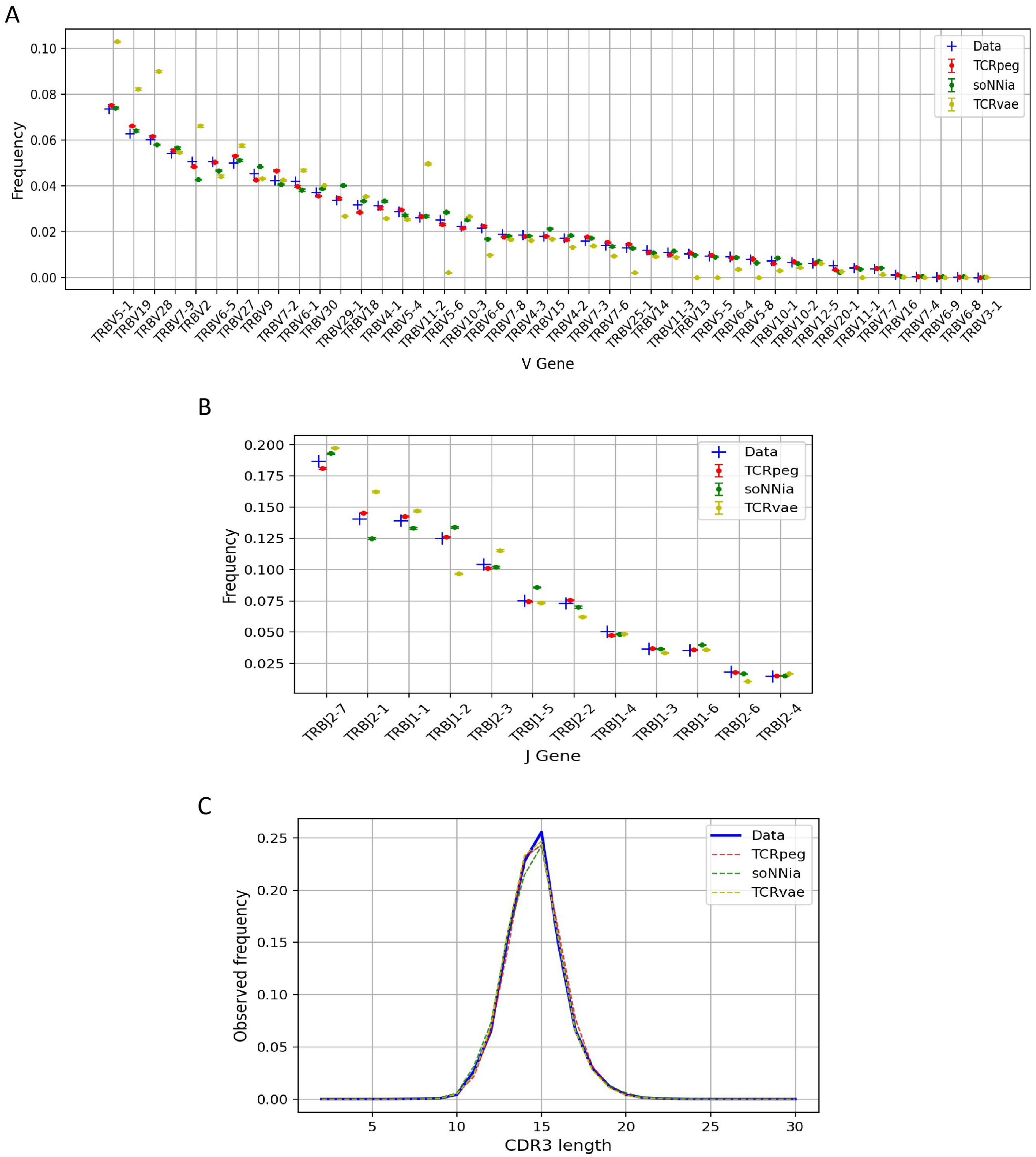


**Figure S9.** Comparisons of (**A**) V gene, (**B**) J gene, and (**C**) length distributions between generated and real sequences when the generative models are inferred from a small subset of the universal TCR pool. Here, we randomly sampled 200 thousand TCRs from the TCR pool and trained each model on this subset. TCRpeg still outperformed the other two models in V and J gene usage distribution with $r \simeq 0.997, 0.998$ compared to $r \simeq 0.993, 0.992$ for soNNia and $r \simeq 0.951, 0.980$ for TCRvae. For length distribution, TCRvae achieved the best performance with $r \simeq 0.992$ compared to $r \simeq 0.990$ for TCRpeg and $r \simeq 0.982$ for soNNia.

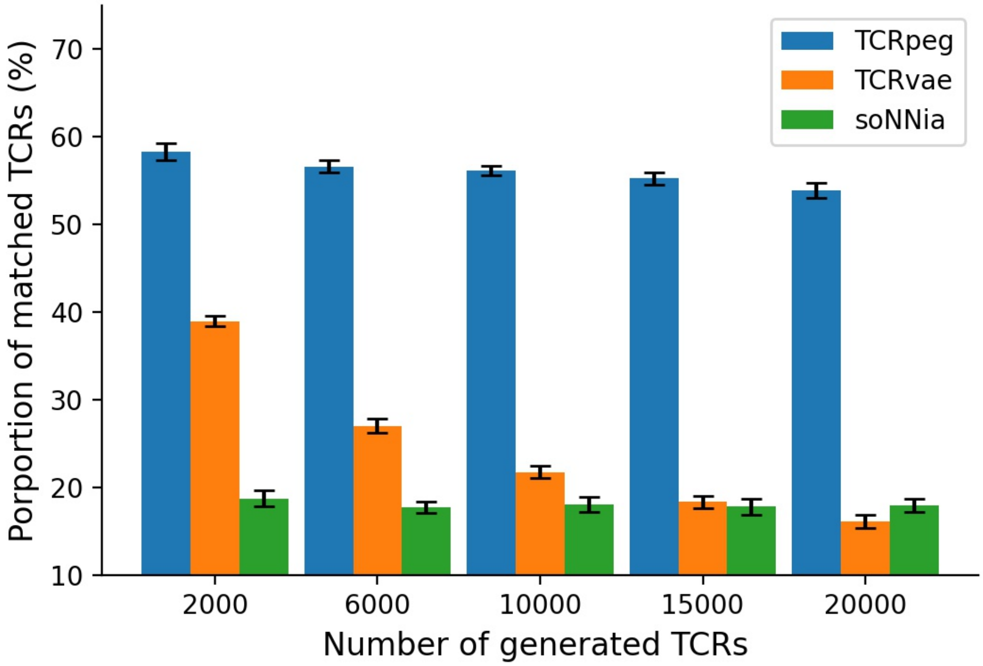


**Figure S10.** Investigation of epitope specificity for TCRpeg-generated TCR sequences using the TCRMatch software. TCRMatch is a software package for predicting TCR specificity based on sequence similarity to previously characterized TCRs. In this experiment, we set the similarity cutoff to 0.9 and assume that those generated TCRs with similarity scores higher than 0.9 to any TCR in the inferring repertoire share the same epitope specificity. Specifically, we inferred the three generative models on the YLQPRTFLL-specific TCRs and used the trained models to generate new TCRs. We observed that more than 50% different numbers of generated TCRs possess epitope specificity to YLQPRTFLL using TCRpeg. These results support our claim that the TCRpeg-generated TCRs may share a hidden similarity to the TCRs used to train the model. In contrast, only less than 40% proportion of the soNNia- and TCRvae- generated sequences possess epitope specificity.

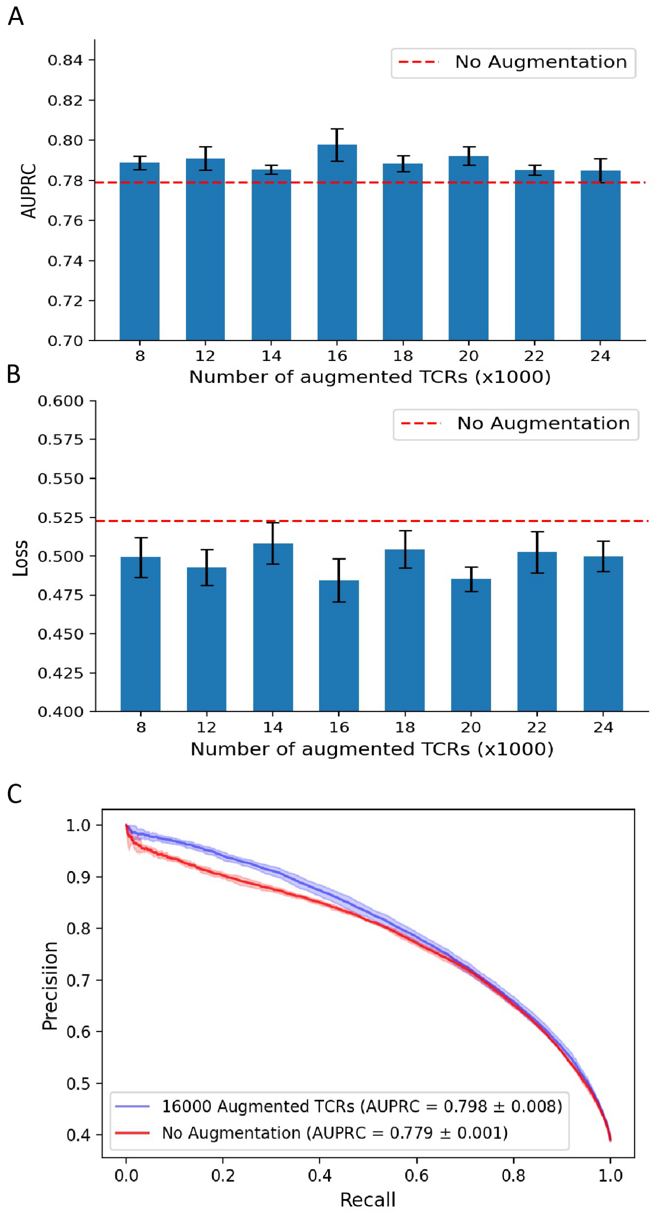


**Figure S11.** Effects of using the TCR-specific data augmentation technique for classifying caTCRs. (**A**) The area under the precision-recall curve (AUPRC) values when data augmentation is applied using the TCRpeg-c model. Within a large range of the number of generated TCR sequences, we observe an enhancement of AUPRC value. (**B**) The binary cross-entropy loss (BCE loss) that is evaluated on the test set. When data augmentation is applied, test loss decreases, which is a positive sign of alleviation of the overfitting problem. (**C**) The precision-recall curve when the number of augmented TCRs is 16,000 where AUC, AUPRC, and the reduction of test loss achieve the highest.

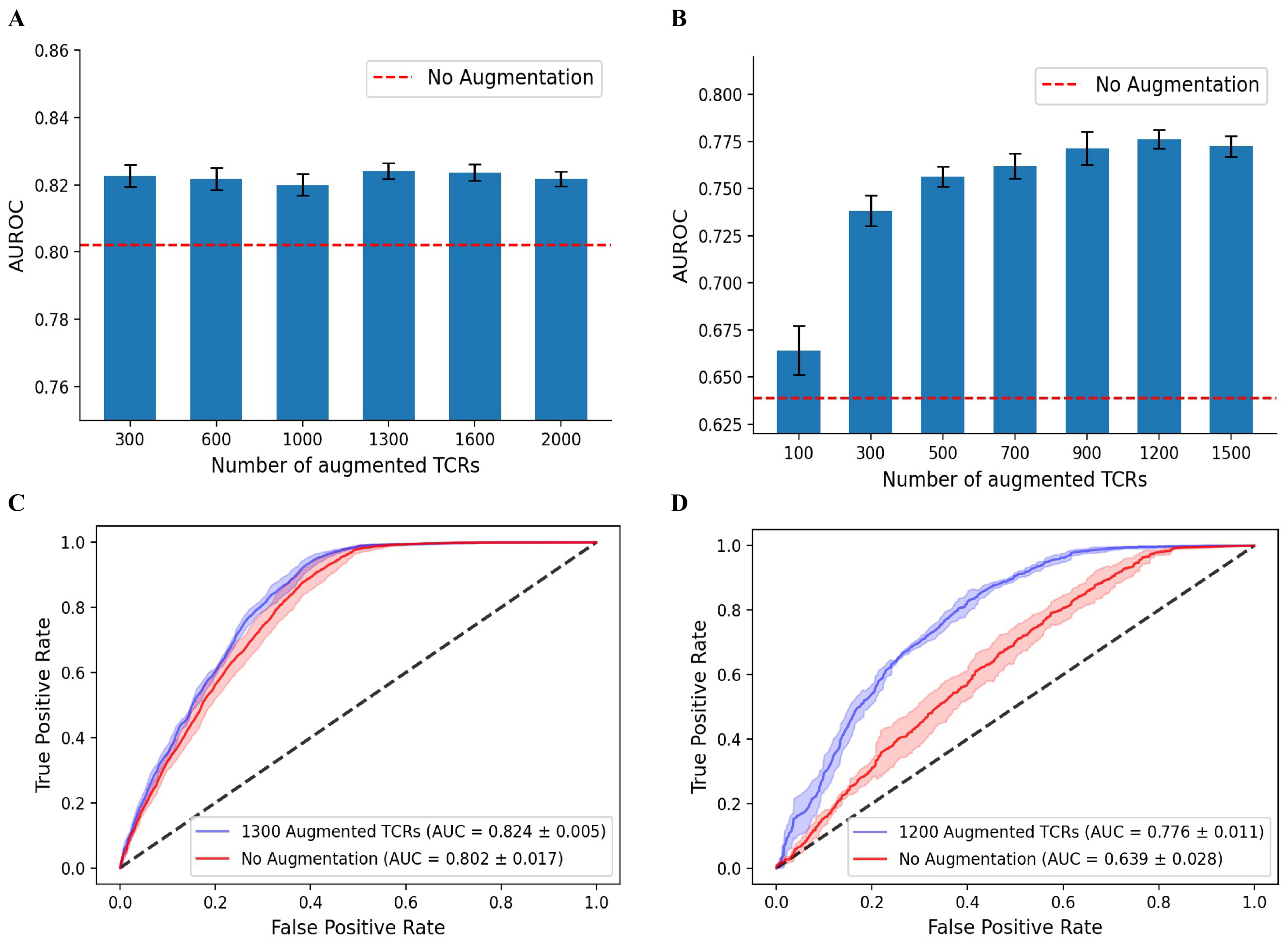


**Figure S12.** Validation of the utility of the TCRpeg-based data augmentation technique using the TCRex model. We collected epitope-specific TCRs for GILGFVFTL and GLCTLVAML from the VDJdb database (downloaded on Mar 21, 2022) with 3406 and 962 positive samples. We randomly sampled the same number of TCRs from the universal TCR pool as the control samples. We trained TCRex in the default setting in all experiments. The TCRpeg model with hidden size and number of layers set to 256 and 3 was trained for 30 epochs and then used to generate new TCRs as augmentation. We show the AUC values when applying data augmentation with a different number of augmented TCRs to the classification of (**A**) GILGFVFTL and (**B**) epitope-specific TCRs of GLCTLVAML. The receiver operating characteristic curves for the GILGFVFTL and GLCTLVAML with the highest performance boost are shown in (**C**) and (**D**) respectively. We observe that applying data augmentation can bring about 2.1% and 21.4% AUC enhancement.
